# The highly diverse and complex plasmid population found in *Escherichia coli* colonising travellers to Laos and their role in antimicrobial resistance gene carriage

**DOI:** 10.1101/2022.08.22.504756

**Authors:** Ann E. Snaith, Steven J. Dunn, Robert A. Moran, Paul N. Newton, David A. B. Dance, Viengmon Davong, Esther Kuenzli, Anu Kantele, Jukka Corander, Alan McNally

## Abstract

Increased colonisation by antimicrobial resistant organisms is closely associated with international travel. This study investigated the diversity of mobile genetic elements involved with antimicrobial resistance (AMR) gene carriage in extended-spectrum beta-lactamase (ESBL) -producing *Escherichia coli* that colonised travellers to Laos. Long-read sequencing was used to reconstruct complete plasmid sequences from 49 isolates obtained from the daily stool samples of 23 travellers over a three-week period. This method revealed a collection of 105 distinct plasmids, 38.1% of which carried AMR genes. The plasmids in this population were diverse, mostly unreported and included 38 replicon types, with F-type plasmids (n=22) the most prevalent amongst those carrying AMR genes.

Fine-scale analysis of all plasmids identified numerous AMR gene contexts and emphasised the importance of IS elements, specifically members of the IS*6*/IS*26* family, in the creation of complex multi-drug resistance regions. We found a concerning convergence of ESBL and colistin resistance determinants, with three plasmids from two different F-type lineages carrying *bla*_CTX-M_ and *mcr* genes. The extensive diversity seen here highlights the worrying probability that stable new vehicles for AMR will evolve in *E. coli* populations that can disseminate internationally through travel networks.

**Impact Statement:** The global spread of AMR is closely associated with international travel. AMR is a severe global concern and has compromised treatment options for many bacterial pathogens, among them pathogens carrying ESBL and colistin resistance genes. Colonising MDR organisms have the potential to cause serious consequences. Infections caused by MDR bacteria are associated with longer hospitalisation, poorer patient outcomes, greater mortality, and higher costs compared to infections with susceptible bacteria.

This study elucidates the numerous different types of plasmids carrying AMR genes in colonising ESBL-producing *E. coli* isolates found in faecal samples from in travellers to Vientiane, Laos. Here we add to known databases of AMR plasmids by adding these MDR plasmids found in Southeast Asia, an area of high AMR prevalence. We characterised novel AMR plasmids including complex ESBL (*bla*_CTX-M_) and colistin (*mcr*) resistance co-carriage plasmids, emphasising the potential exposure of travellers to Laos to a wide variety of mobile genetic elements that may facilitate global AMR spread. This in-depth study has revealed further detail of the numerous factors that may influence AMR transfer, therefore potential routes of AMR spread internationally, and is a step towards finding methods to combat AMR spread.

**Data Summary:** Long-read sequencing data is available through National Center for Biotechnology Information under the BioProject PRJNA853172. Complete plasmid sequences have been uploaded to GenBank with accession numbers in supplementary S1. The authors confirm all supporting data, code and protocols have been provided within the article or through supplementary data files.

## Introduction

Infections caused by antimicrobial-resistant organisms are harder to treat, lengthen hospital stays, increase mortality rates, and place a significant financial burden on healthcare institutes (1). It is increasingly important to characterise the mechanisms that allow antimicrobial resistance (AMR) to spread worldwide, compromising treatment options for many bacterial pathogens. The rapid spread of AMR has been closely associated with international travel (2–5). Clinically relevant AMR determinants are commonly found in Gram-negative bacteria colonising travellers returning from regions with highest AMR prevalence, including South-East Asia (3, 6, 7).

*Escherichia coli*, a Gram-negative human gut commensal and opportunistic extraintestinal pathogen, is an important vector for AMR (8, 9). As exemplified by pandemic multi-drug resistant (MDR) lineages such as ST131 (10, 11), *E. coli* is capable of acquiring and maintaining multiple AMR determinants and exhibiting resistance to multiple classes of antibiotics. The presence of combinations of AMR genes can significantly impact therapeutic options. The limited treatment options for ESBL-resistant organisms make these a cause of concern. Colistin is a last-resort antibiotic included in the Reserve category of the WHO Essential Medicines List (12). In *E. coli* and other Gram-negative bacteria, carriage of both colistin resistance genes and production of extended spectrum beta-lactamases (ESBLs) is concerning as it suggests a potentially stable environment for the accumulation of further resistance, for example, carbapenems, severely limiting treatment options (9, 13, 14). ESBL resistance genes, including *bla*_CTX-M_ variants, and *mcr* genes that confer colistin resistance are commonly found in *E. coli* carried by returning travellers (3, 6, 7, 15). AMR genes can move intra- or inter-cellularly and accumulate at single sites in association with mobile genetic elements (MGEs) (16–19). Plasmids are extrachromosomal genetic elements that can transfer horizontally between bacteria of the same or different species and are strongly associated with the spread of AMR (20, 21). In *E. coli* and other members of the Enterobacterales, AMR genes have been found in many different plasmid types (20). Reports of epidemic and internationally-distributed plasmids (22–24), highlight the threat successful plasmid lineages pose and the importance of understanding the mechanisms by which they acquire and accumulate AMR genes.

We recently characterised the dynamics of acquisition of multi-drug resistant Gram-negative organisms in real time during travel to Vientiane, Laos. These MDR Gram-negative organisms had a surprisingly high co-prevalence of ESBLs and colistin resistance genes (7). Here we explore the pattern of AMR carriage and context in *E. coli* from that cohort using long-read sequencing to understand the contexts of AMR genes and the role of MGEs, particularly plasmids, in the acquisition of drug-resistant *E. coli* by travellers to a region of high AMR prevalence. The 49 representative isolates selected for long-read sequencing in this study were collected on a daily basis in an area of high AMR prevalence, enabling continuous monitoring of the drug-resistant *E. coli* that colonised study participants. Continuous sampling facilitated the examination of common and circulating plasmids in the *E. coli* population, across a variety of sequence types (STs) and at different study time points. Our data shows the diversity and widespread distribution of numerous distinct AMR plasmids acquired by these travellers in Vientiane, Laos and the multiple different potential routes of AMR spread by plasmids and highlighted the complex nature of plasmids carrying both ESBL resistance and *mcr* genes.

## Methods

### Study design and Sample source

The *E. coli* isolates used here were collected as part of a study (7) looking at the dynamics of gut colonisation of 21 volunteers attending a medical course in Vientiane, Laos. Faecal samples were taken daily during the 22-day period. Samples were processed, shipped, stored and handled as previously described in Kantele et al (7). Here ESBL-positive isolates were cultured from faecal samples after initial screening on CHROMagar ESBL agar plates (CHROMagar, Paris, France) at the Microbiology Laboratory of Mahosot Hospital, Vientiane, Laos and after transportation further screened with chromID ESBL chromogenic medium (bioMérieux) by University of Helsinki, Helsinki, Finland (7). These ESBL isolates (7) were used in this study. In order to explore the pattern of AMR in this traveller dataset and the role of MGEs we prepared hybrid assemblies and annotated plasmid sequences identified to locate resistance genes and potential routes for spread.

### Selection of *E. coli* isolates for MinION Sequencing

Whole genome sequences of 306 ESBL-positive Gram-negative isolates from the Laos study were previously generated (7) and are available under NCBI BioProject accession number PRJNA558187. Illumina-generated whole genome sequences of *E. coli* isolates (n=219) were sequence typed using mlst (v2.15) (https://github.com/tseemann/mlst). Isolates were selected for MinION sequencing (n=49) by deduplication of longitudinal patient samples, and by using abricate (v.0.8.10) (https://github.com/tseemann/abricate) to identify unique elements such as plasmid replicons (plasmidfinder (v.0.8.10)) and antibiotic resistance genes (resfinder (v.0.8.10)) (36) from the short read assemblies generated with SPAdes (v. 3.13.0).

### DNA Extraction and Sequencing

*E. coli* were cultured overnight on UTI Chromogenic agar (Sigma) at 37°C. After purity checks single colonies were subcultured overnight in LB Broth (Miller) (shaking, 37 °C). For the majority of isolates DNA was extracted using Monarch Genomic DNA Extraction kit (NEB), but in some instances a lower quantity and quality yield was obtained. In these instances, we noticed atypical precipitates and opted for extraction using phenol chloroform with Cetyltrimethylammonium bromide (CTAB). The extracted DNA was sequenced over 4 runs MinION (Oxford Nanopore Technology), using R9.4.1 flow cells. Three runs were prepared using Ligation Protocol (LSK-SQK109), and one run with the Rapid Barcoding Sequencing Kit (SQK-RBK004), with both protocols modified to a one-pot implementation.

### Long read sequencing analysis

Data was basecalled using Guppy (v0.5.1) https://github.com/nanoporetech/pyguppyclient). The quality of the data was examined using NanoPlot (v1.28.2) (25). In read files where coverage was high, filtlong (>100X) (v0.2.0) (https://github.com/rrwick/Filtlong) was used to select the best-read files available and the coverage was limited to 100X. Barcodes were trimmed using qcat (v1.0.1) (https://github.com/nanoporetech/qcat). Hybrid assemblies using the Illumina reads were created with Unicycler (v0.4.7) (26) and visualised in Bandage (v.0.8.1) (27). In some instances, Unicycler was unable to fully resolve the assembly; in these cases we took a long-read first approach, using Flye (v2.6) (28) to assemble the long read set into a gfa file that was then provided to the Unicycler pipeline. Assemblies were analysed using abricate (v.0.8.10) with the resfinder (29, 30) and plasmidfinder databases (31). TigSPLIT (https://github.com/stevenjdunn/TigSPLIT) was used to extract contigs, allowing detailed analysis of the location of plasmid replicons in relation to resistance genes present in each isolate.

### Analysis of plasmids sequences

Plasmid contigs were annotated using Prokka (v1.14.6) (32). abricate (v.0.8.10) was used to identify plasmid replicons with the PlasmidFinder database and AMR genes were identified with Resfinder (85% coverage and identity cut offs). In situations where plasmidfinder did not provide a type, plasmid contigs were compared to known plasmid replicons for type assignments. Where possible, plasmid replicons were sub-typed using the PubMLST database (https://pubmlst.org/) (33, 34). Snapgene (v5.2.4) was used to visualise Prokka-annotated plasmid contigs. ISFinder (35) was used to identified insertion sequence (IS) elements. NCBI BLAST (v2.5.0+) was used to compare plasmids to reference plasmids (Supplementary S2). The NCBI non-redundant nucleotide database was queried with BLASTn to determine whether plasmids identified here had been seen elsewhere. BLASTn and tBLASTn were used to compare similar plasmid regions and to identify homologs of known plasmid genes. Mashtree (v1.2.0) (36) and Panaroo (v1.2.3) (37) were used to compare plasmids. ISEScan (v.1.7.2.3) was used to quantify IS elements in plasmid assemblies at an element-family level (38).

### Identification of small plasmids from Illumina dataset

NCBI BLAST (v2.5.0+) was used to search draft genomes for plasmid-specific 100 bp signature sequences (Supplementary S3), as described previously (22). This facilitated the detection of specific plasmid lineages whether they were represented by single or multiple contigs in the Illumina dataset (7).

## Results

### Isolates from Laos contain a broad diversity of plasmids and resistance genes

A total of 163 complete plasmids were obtained from the hybrid-assembled genomes of 49 *E. coli* isolates that colonised 21 participants and their contacts (Supplementary S4). Isolates harboured 1 to 8 plasmids each, except for a single ST1722 isolate that did not contain any plasmids. The 163 plasmids were typed by size, replicon type and AMR gene carriage, and identical or almost-identical plasmids (Table S3) were deduplicated, resulting in a total of 105 distinct plasmids. These 105 plasmids were named pLAO1-pLAO105 (Supplementary S1 and S4) and ranged in size from 1,531 – 259,739 bp, with around half (53.3%, n=56) smaller than 25 kb. Forty of the 105 plasmids (38.1%) contained one or more AMR genes. Plasmids that did not carry AMR genes were generally smaller (mean size: 23,554 bp), than those that did (mean size: 97,260 bp). However, pLAO10 (GenBank accession OP242224), pLAO78 (OP242289) and pLAO84 (OP242239) are notable ColE1-like plasmids that carry 1-5 AMR genes each and range in size from just 5,540 bp to 22,368 bp (Table S4).

Thirty-eight different replicon types were identified in the collection of 105 plasmids. Types present amongst small plasmids included: theta-replicating plasmids that initiate replication with RNA primers (θ-RNA, n=26) or replication proteins (θ-Rep, n=9), rolling-circle plasmids (n=6), and Q-type plasmids that replicate by strand-displacement (n=3). Amongst large plasmids (>25 kb), F-types (n=22) were dominant, and their replicons were sub-typed using the PubMLST database (Supplementary S4). Other large plasmids include phage-plasmids (n=9, including Y or p0111 types) and X-types (n=5). Nine large plasmids were co-integrates, which included multiple replicons of disparate types and all contained one or more AMR genes (Figure 1). Ten plasmids could not be assigned a type (Figure 1). In total, 76.3% (n=29) of plasmid types contained AMR genes, with the most prevalent being FII-46:FIB-20 (n=4), X1 (n=3), θ-RNA (n=3), FII-2 (n=2), FII-46:FIB-10 (n=2), p0111 (n=2) and Y (n=2) (Figure 1, Supplementary S4). AMR plasmids were present across all participants throughout the study (Supplementary S4 & S5), with carriage occurring sporadically and transiently during the study period.

**Figure 1.**
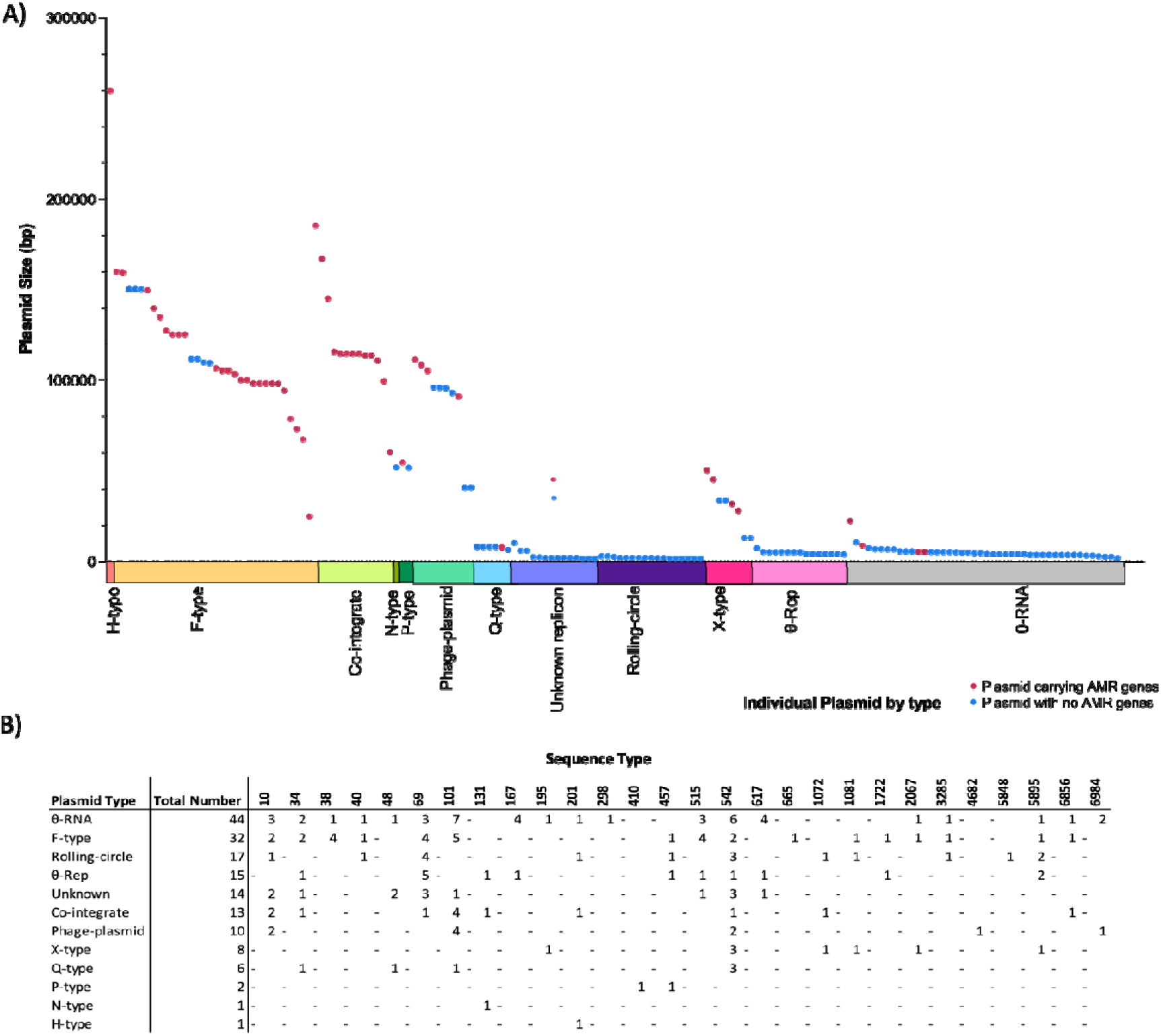
An overview of plasmid diversity showing plasmid size and AMR gene carriage (A) and plasmid type within sequence type (B) for all plasmids identified

We found evidence suggesting the circulation of plasmids within Lao *E. coli* in the traveller population. One example, the ColE1-like plasmid pLAO84 (GenBank accession: OP242239), which carries the tetracycline resistance determinant *tet*(A), was found in the complete genomes of *E. coli* of two different STs, ST195 and ST34, that were acquired by two different participants (Pt33 and Pt40). Mapping Illumina reads against the complete pLAO84 sequence confirmed plasmid circulation, with its complete or fragmented sequence detected in seven additional *E. coli* isolates (Supplementary S6). These isolates were obtained from six different participants (including Pt33 and Pt40) over a 10-day period. Another example of a potential circulating plasmid, the Q1 plasmid pLAO60 (GenBank accession OP242237) that carries aminoglycoside and beta-lactam resistance genes (Table S4), was present in the complete genome of the ST542 isolate (LA124) and in one genome (LA230) in the wider Illumina dataset (Supplementary S4 & S7). The LA230 isolate Q1 plasmid was missing a 101 bp segment that was likely lost in a homologous recombination event.

Another notable finding was the presence of phage and virulence plasmids in this collection. A Y-type phage-plasmid, pLAO59 (GenBank accession OP242238), appears to have lost key genes for phage body synthesis, potentially in deletion events mediated by IS*Kpn26* and IS*1294*, and instead carries genes that confer resistance to six antibiotic classes and genes that confer resistance to mercury, copper, and silver (Supplementary S8). The FII-18:FIB-1 pLAO32 is related to the colicin V (ColV) virulence-resistance plasmid pCERC3 (39) and contains virulence genes, including those for the aerobactin and Sit siderophore systems, in addition to multiple drug resistance genes (Supplementary S8), but lacks the genes for ColV.

### Variation within individual plasmid types, with diverse and complex resistance regions

In addition to the variety of different plasmid types, long-read sequence data allowed us to observe diversity amongst plasmids of the same type (Supplementary S4-S9). There was considerable genetic diversity in plasmids carrying the FII-2 replicon (Figure 2). FII-2 plasmids pLAO44 (GenBank accession OP242233) and pLAO37 (OP242229) in ST69 and ST101 strains only contained FII-2 replicons, while FII-2 plasmid pLAO82 (OP242230) in an ST34 isolate carried an additional θ -RNA replicon and plasmids pLAO100 (ST40, OP242240) and pLAO103 (ST457, OP242243) carried an additional FIB-10 replicon (Figure 2 & 3). Diversity also occurred in the resistance regions of these plasmids (Figure 2 & Supplementary S9). FII-2 plasmids carried multiple resistance genes, including various combinations of *bla*_CTX-M-27_, *bla*_CTX-M-55_ and *mcr-3.4* (Figure 2 & 3). Plasmid types FII-2 (pLAO37), FII-2:θ-RNA (pLAO82) and FII-46:FIB-like (pLAO69, OP242232) incorporated resistance regions with an accumulation of multiple resistance genes and co-carriage of *mcr* and *bla*_CTX-M_ genes (Figure 3B). FII-2:θ-RNA, pLAO82, demonstrated even more complexity with further accumulation of resistance genes as it also carried an additional *mcr-3.4* located between copies of IS*26* and IS*Kpn40* (Figure 3). There is widespread global distribution of the resistance regions found in these FII-2 plasmids which have been identified in multiple countries, some of which are not close to Laos (Supplementary S10). We found no link between the ST of the isolate and the type of plasmid AMR genes carried by that ST (Figure 3A).

**Figure 2.**
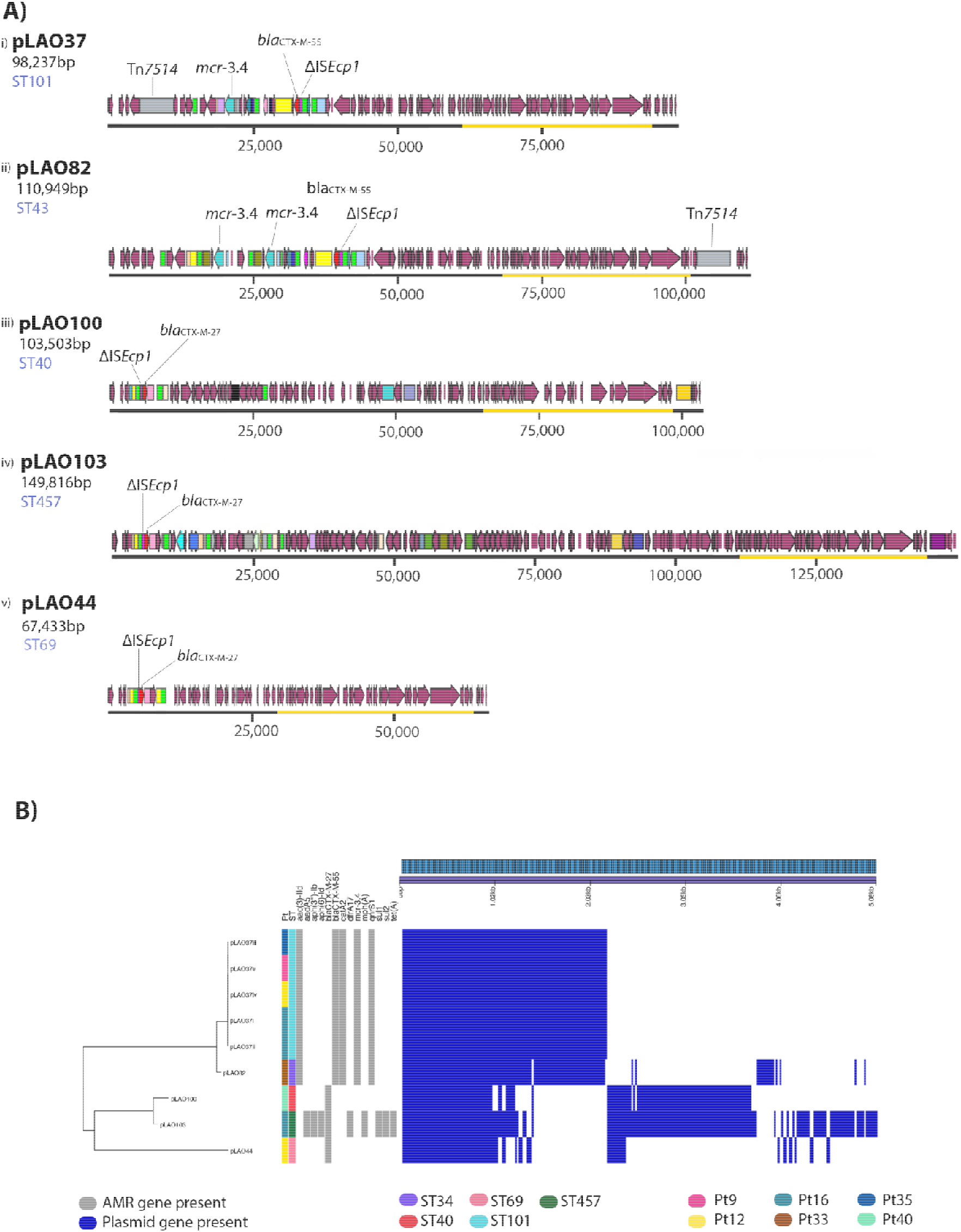
Diversity within FII-2 plasmid replicon group showing variable plasmid sizes and resistance genes. **A) Annotated FII-2 plasmid group maps indicate the location of *bla_C_*_TX-M_ and *mcr* genes.** Maroon arrows represent Prokka-annotated genes. All other brightly coloured arrows represent antibiotic resistance genes. Transposable elements are displayed as brightly coloured boxes. Notable elements are IS*2*6 (bright green), IS*Ecp1* (bright pink), Tn*1721* (pale blue) and IS*Kpn26* (lavender). The plasmid transfer (*tra*) region is highlighted with an orange outline. **B) IQ-tree of FII-2 plasmids core alignment showing AMR genes and gene/presence absence matrix**. The participant (Pt) and ST of the isolate from which the plasmid was identified are displayed in different colours. Resistance genes identified using Abricate (grey = present) are shown alongside the gene presence/absence profile from Panaroo (royal blue = present).

**Figure 3.**
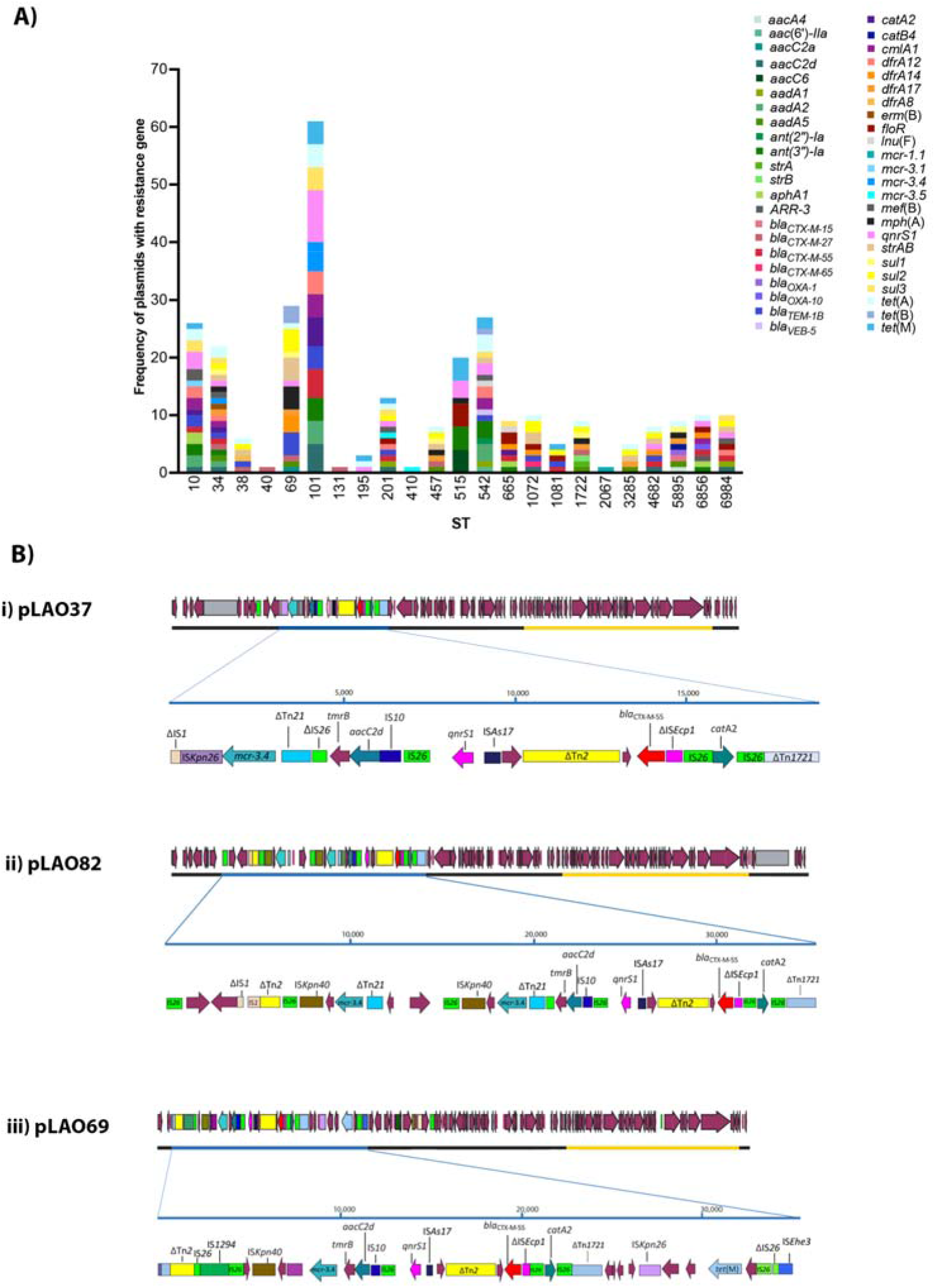
Antimicrobial resistance gene carriage diversity by A) ST and B) within *mcr* and *bla*_CTX-M_ co-carriage plasmids. All maroon genes are prokka annotated genes. All other brightly coloured genes are antibiotic resistance genes. Transposable elements (e.g. transposons, insertion sequences) are displayed as brightly coloured boxes). The plasmid transfer (*tra*) region is highlighted with an orange outline.

### Co-carriage of *bla*_CTX-M_ and *mcr* genes occurs in multiple resistance region configurations

Three plasmids in this collection carried both *bla*_CTX-M_ and *mcr* genes (Figure 3B). All three *bla*_CTX-M_/*mcr* co-carriage plasmids were F-types, but they differed in size and plasmid replicon type. The 98,237 bp plasmid, pLAO37, carried only an FII-2 replicon and was found in five ST101 *E. coli* isolates. The 110,949 bp pLAO82, was a multi-replicon co-integrate plasmid carrying both FII-2 and θ-RNA replicons and was found in one ST34 *E. coli* isolate. The 105,425 bp plasmid, pLAO69 carried FII-46 and FIB-20 replicons, and was found in two ST10 *E. coli* isolates. All three of these plasmids have typical F-type plasmid backbones that include genes for replication, stable maintenance, conjugative transfer and establishment in new hosts (39, 40). Each plasmid contained a complete and uninterrupted transfer region, suggesting all three have the capacity for self-mediated conjugation (41).

Detailed comparison of the FII-2 plasmids pLAO37 and pLAO82 showed that the backbones are almost identical apart from a recombination patch (approximately 7 kb). pLAO37 and pLAO82 have the same FII-2 *repA1* gene. The AMR genes in both pLAO37 and pLAO82, are located in a complex region bounded by IS*26* at the left end and Tn*1721* at the right end (Figure 3B). These regions are comprised of sequences from multiple mobile genetic elements with distinct origins and include genes that confer resistance to beta-lactams (*bla*_CTX-M_); colistin (*mcr-3.4*); aminoglycosides (*aacC2d*); chloramphenicol (*catA1*) and quinolones (*qnrS1*). An additional θ-RNA replicon in pLAO82 is part of a small plasmid that has been captured and incorporated into the resistance region. pLAO37 and pLAO82 contain an extra partial Tn*21* and partial IS*26* region inserted in between *tmrB* and *mcr* gene (Figure 3B). The pLAO82 resistance region also includes an additional copy of *mcr-3.4*. The *mcr-3.4* in pLAO37 was truncated by IS*Kpn26*, which removed the terminal 32 bp of the gene. This configuration of IS*Kpn26* and *mcr-3.4* is not present in any other sequence deposited in the GenBank non-redundant nucleotide database. A novel transposon, Tn*7514*, was present in both pLAO37 and pLAO82 but has inserted in two different backbone locations (Figure 2B, Supplementary S11 & S12)(42).

Although largely syntenic, the nucleotide identity of the pLAO69 backbone differs significantly from that of pLAO37 and pLAO82, consistent with its distinct replicon type. The FII-46 *repA1* gene of pLAO69 is only 94% identical to FII-2 *repA1* of pLAO37 and pLAO82. The pLAO69 transfer region also differed significantly matching only 89% of pLAO37 and pLAO82 transfer region with an overall identity of 97.2% in a BLASTn comparison. pLAO69 carried *mcr-3.1*, which differs from the *mcr-3.4* found in pLAO37 and pLAO82 (Figure 3B). However, inspection of the resistance region in pLAO69 that contains *mcr-3.1* revealed that it is largely comprised of the same confluence of mobile elements found in the resistance regions of pLAO37 and pLAO82 (Figure 3B). In addition to their identities, the configuration of these elements was the same in pLAO69 and pLAO37/pLAO82.

### Insertion sequence type, abundance and diversity is associated with AMR gene carriage

The vast majority (n=36) of the 40 AMR plasmids contained one or more IS. The four exceptions were plasmids that contained AMR genes found in association with gene cassettes (*bla*_VEB_ in pLAO60 and *mcr-1.1* in pLAO41 and *aadA2*, *ant(3″)-Ia*, *cmlA1*, *dfrA12* in pLAO78), or Miniature Inverted-repeat Tandem Elements (MITEs) (*tet*(A) in pLAO84). AMR plasmids carried almost three times more unique families of IS than their non-AMR counterparts (Figure 4A), with an average of 6.75 unique IS families in AMR plasmids (range 1-11), vs 2.4 in non-AMR plasmids (range 1-8).

**Figure 4.**
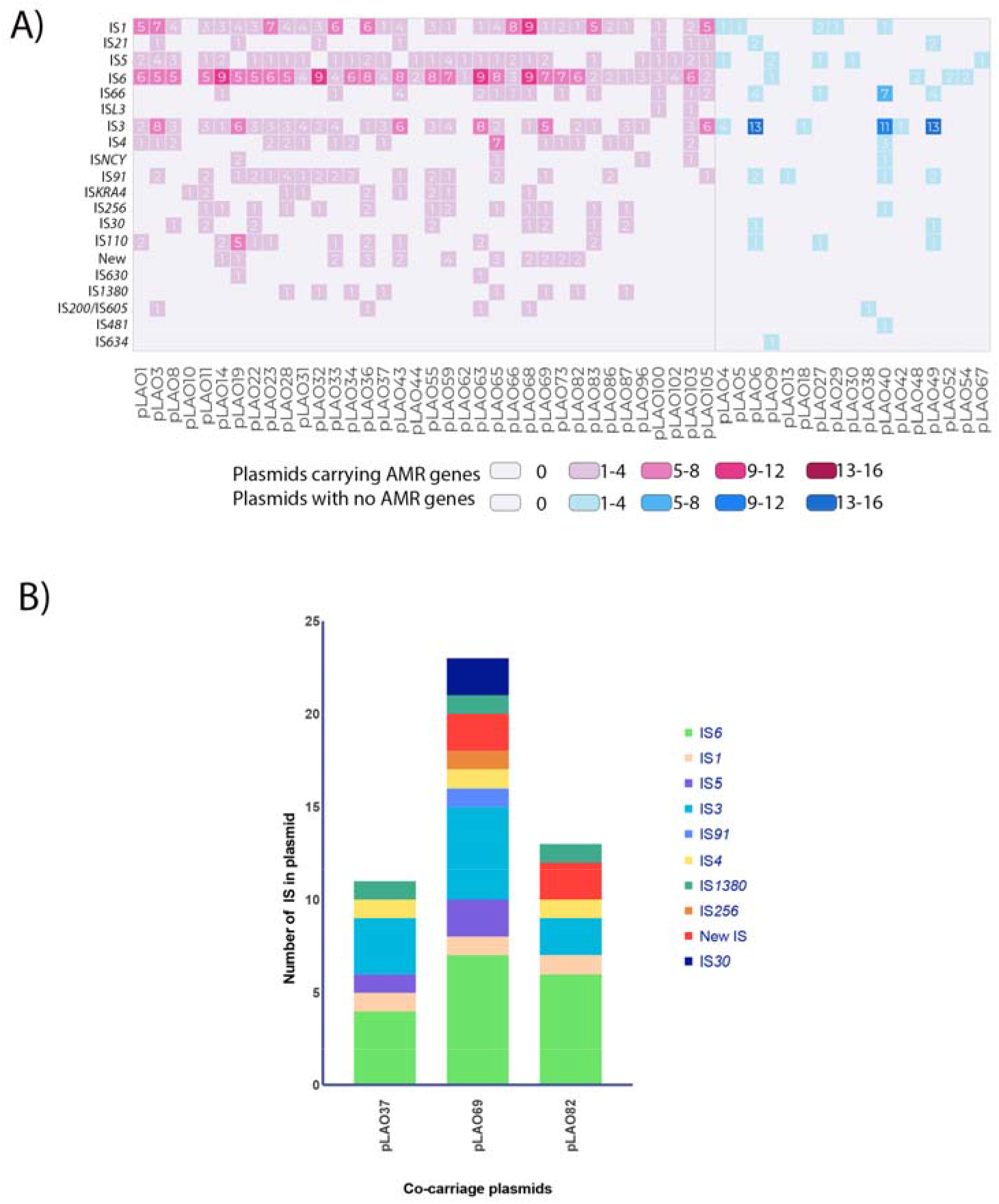
IS families present in A) all plasmid types displaying AMR gene presence/absence and IS linked to B) *mcr* and *bla*_CTX-M_ co-carriage plasmids

AMR plasmids contained an abundance of IS*6*/26 and IS*1*-family elements. Of the 36 AMR plasmids that did carry IS elements, all but one carried at least one IS*6*-family element (Figure 4A), and IS*1* family was present in 83% (n=30). IS*3* and IS*5* were also common, present in 75% (n=27) and 78% (n=28) of AMR plasmids, respectively. Differences in IS*6* family element carriage were seen between AMR and non-AMR plasmids, with AMR plasmids containing a mean of five IS*6* family elements per plasmid (range 0-9). In contrast, only 24% (n=4) of non-AMR plasmids carried IS*6* family elements, with only 1-2 IS*6* elements per plasmid (Figure 4A). IS6-family elements were present in all plasmids that co-carry *bla*_CTX-M_ and *mcr* genes (Figure 4B), where they were the most prevalent IS family. There was no clear association between IS families and co-carriage of the *mcr* and *bla*_CTX-M_ genes (Figure 4B). In plasmids where there was co-carriage of *mcr* and *bla*_CTX-M_ genes, IS*6* and IS*3* were the most abundant IS families, with all co-carriage plasmids additionally carrying IS*1380*, IS*1* and IS*4*. In each co-carriage plasmid, more than six different IS families were identified, and of the IS identified between 30%-46% were IS*6* family (Figure 4B) with IS*26* the most prominent element.

## Discussion

In this study we conducted an in-depth investigation of the role of plasmids in the alarmingly high levels of AMR found in *E. coli* that colonised the GI tract of travellers to Laos (7). We have revealed an enormous diversity in the plasmids of this *E. coli* population, particularly in their resistance regions, many of which contained multiple AMR genes. Concerningly, the majority of the 40 AMR plasmids (n=30, 75%) contained ESBL genes, a colistin resistance gene or both. Our data showed the abundance and importance of plasmid types F-type, X-type, Q-type and ColE1-like plasmids as vectors for AMR gene spread in *E. coli* in Laos. AMR plasmids accounted for 38.1% (n=40) of the 105 distinct plasmids, 17 of these 40 AMR plasmids identified were F-type (42.5%) highlighting the importance of F-type plasmids as carriers of AMR genes (including ESBLs, *mcr*). F-type plasmids are known to carry AMR genes (43) and are an important factor in the high incidence of AMR carriage in this study. Although four types dominated, the AMR plasmid population identified in travellers to this region of Laos was extremely diverse with 29 different plasmid types, including phage and virulence plasmids, found to carry AMR genes. Both AMR and non-AMR plasmids were identified throughout the study, including several from baseline faecal samples (7) where plasmids may have been acquired from travel to Laos or potentially from other travel.

A variety of less anticipated vehicles for AMR genes were identified in these Lao *E. coli*. Small ColE1-like θ-RNA plasmids are common in *E. coli* (21, 44) and their high prevalence would be expected. These θ-RNA (n=26) were the most prevalent plasmid in the collection, followed by F-types (n=23) (Figure 1) but in contrast to F-type plasmids, only a small number of θ-RNA plasmids (n=3) carried AMR genes. This is consistent with previous findings that ColE1-like plasmids occasionally carry resistance genes, including aminoglycoside and beta-lactam resistance determinants (45). While relatively uncommon, the importance of small, high-copy number plasmids to AMR should not be underestimated, as these can serve as platforms for the evolution of new resistance phenotypes (45, 46). The *tet*(A)-carrying ColE1-like plasmid pLAO84 clearly demonstrates the capacity these small plasmids have for disseminating resistance genes. pLAO84 was found in multiple *E. coli* STs in this collection, suggesting that it was circulating in the local *E. coli* population before being acquired by multiple study participants (Supplementary S6). Additionally, the presence of a plasmid almost identical to pLAO84 in GenBank (GenBank accession CP057097.1) indicates this ColE1-like plasmid lineage has already spread internationally, as it was present in *Escherichia fergusonii* isolated from pig faeces in the United Kingdom.

A Y-type phage-plasmid (pLAO59, OP242238, Supplementary S8) highlighted an additional opportunity for AMR spread since there is a chance for co-selection of this MDR plasmid due to carriage of metal resistance genes e.g. for silver and copper. pLAO59 appeared not only to have lost key phage genes required for the lytic lifestyle (47), apparently as a result of deletions by insertion sequences, but carried multiple AMR genes and heavy metal resistance genes. Co-resistance and cross-resistance can cause co-selection of bacteria carrying metal resistance and AMR genes, with the metal resistance gene causing maintenance of the AMR gene (48, 49). In Vientiane, Laos it has been reported that environmental samples sourced near municipal solid waste landfill showed heavy contamination with heavy metals including copper at levels higher than WHO permissible standards (50), indicating the possibility that co-selection by metal resistance genes is a real environmental pressure in the Vientiane area from which these isolates were collected. Silver and copper can co-select for various AMR genes in *E. coli* including tetracycline and sulphonamide resistance genes, which were also found in pLAO59 (51, 52).

Multiple combinations of AMR resistance genes were found in multiple genetic contexts associated with various mobile genetic elements. *mcr* genes for example were found next to multiple TEs (Figure 3, Supplementary S4) and mainly on F-type plasmids, consistent with literature indicating that *mcr-3.4* is associated with the FII-type (53, 54). IS*Ecp1* is known to contribute to the spread of *bla*_CTX-M_ (55), but interestingly only one complete IS*Ecp1* was identified (pLAO32) as part of a *bla*_CTX-M-55_-containing transposition unit flanked by the 5 bp target site duplication TAACA. Most *bla*_CTX-M_ genes in this collection were associated with complete copies of IS*26* and partial IS*Ecp*1 (Figure 2, 3, Figure 4A and S8-S9). IS*26* from the IS*6/26* family is known to play a key role in AMR gene dissemination (56–59). IS*26* carriage predisposes plasmids to insertion of additional IS*26* and any associated AMR genes, which facilitates the accumulation of AMR genes at single sites leading to multi-drug resistance phenotypes (56, 60). This appears to be the case with pLAO37, pLAO82 and pLAO69 featuring IS*26* and based on what is known of IS*26* behaviour (56), IS*26* is likely to have played an important role in the assembly of their complex co-carriage resistance regions (Figure 3).

Analysis of plasmid contigs from 49 *E. coli* hybrid assemblies has confirmed the vast and diverse genetic context of AMR in the Kantele et al dataset isolated in Laos, and highlighted that as well as multiple unique colonising strains (7), there are multiple distinct plasmids present in this dataset. Long-read sequencing has been critical in highlighting the key role insertion sequence elements may play in the formation of this complex set of MDR plasmids that create a risk of spread AMR. This previously unreported cohort of MDR plasmids offers the alarming prospect that one of these will create a stable configuration for the creation of a successful pandemic MDR plasmid.

## Author Statements

### Author contributions

AM and JC conceived the idea for this analysis. AK, EK, DABD, PNP, VD collected samples, isolated the strains and provided the isolates and epidemiological data in Laos. AS and SD performed long-read genomic sequencing and sequence processing. AS, SD and RM performed genomic analysis, data interpretation and data visualisation. AS, RM and SD drafted the article. AM discussed results and edited draft. JC, AK, PN and DD critically reviewed the draft. All authors read and approved the final manuscript.

### Ethical Approval

The study protocol (see Appendix 1 of reference (7)) was approved by the Ethics Committee of the Helsinki University Hospital, Helsinki, Finland, and the Ethics Committee of Northwest and Central Switzerland, Basel, Switzerland. All participants provided written informed consent.

## Acknowledgements

We are very grateful to the volunteers and the staff of the Microbiology Laboratory, Mahosot Hospital, Vientiane, Laos, who processed the samples and for the support of the directors and staff of Mahosot Hospital and the Ministry of Health, Lao PDR.

