## Supplementary figures and tables for "The highly diverse and complex plasmid population found in *Escherichia coli* colonising travellers to Laos and their role in antimicrobial resistance gene carriage"

**Supplementary Data**

**Supplementary S1** – In separate excel document.

| **Plasmid Type** | **Name** | **Genbank Accession number** |
| --- | --- | --- |
| FII-2 | p39R861-3 | MK092064.1 |
| IncX1 | pSRC22-2 | MN256104 |
| IncQ | RSF1010 | M28829 |
| FII-18 | pCERC3 | KR827684 |
| IncY | P1 | AF234172 |
| ColRNAI | pCERC7 | KX356458 |

**Supplementary S2** – Reference plasmids used to annotate key genes on plasmid types identified in this dataset

| **Section Number** | **Section looked for in Illumina dataset** |
| --- | --- |
| ColE1-like plasmid (pLAO84) MITESen1 to backbone section 1 | TTAAATAATGCGCTTAACGTACAAAAAATTCCGATCTCCAAACTGACCCCTTCTTCGCCTCTTATGTTTCTCGAATGAATCGATGGGGACAGGAAATGTT |
| ColE1-like plasmid (pLAO84) MITESen1 to backbone section 2 | ATTTATTTTTCATGAAGTTGCGATAAAAATCGCAGCTGCGTTAGGTGTATGGGGTCAGTTTGGATATAGAGAATTATTGTACGGTAAGCCTGTTTTTTGA |
| IncQ1 Section from pLAO60 | CCGTTAACTGTCACGCCCCCCCGTTAACTGTCACGAACCCCCCGTTAACTGTCACGCCCCCCGTTAACTGTCACGAACCCCCCGTTAACTGTCACGCCCCC |

**Supplementary S3** – DNA sections searched for in the Illumina dataset (6) to look for circulating ColE1-like plasmid (pLAO84) and IncQ1 plasmid (pLAO60)

**Supplementary S4**– In separate excel document.

**Supplementary S5** - In separate excel document

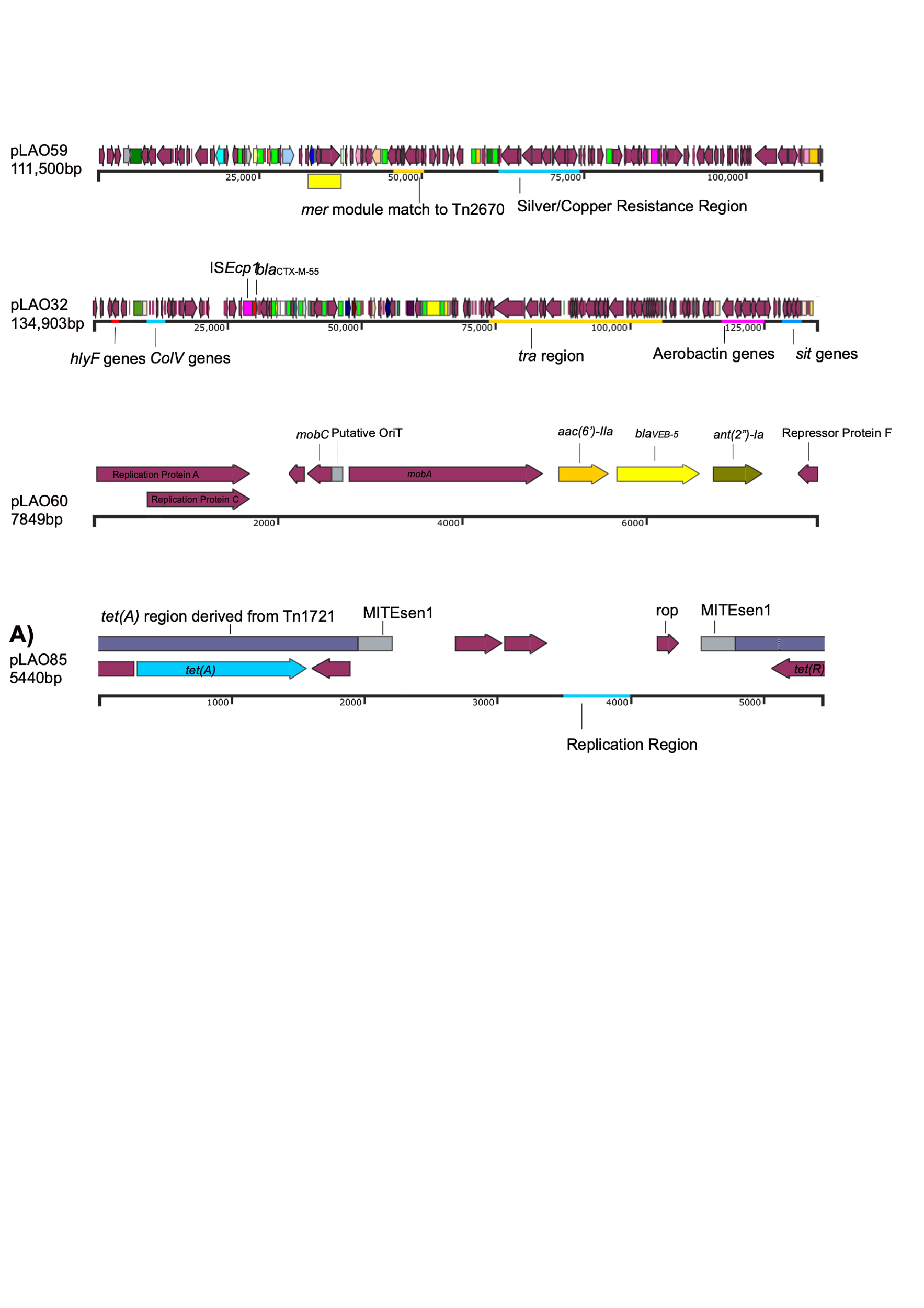

**B)**

| **MITSen1 and MITSen2 Blast Contig Hit** | **Findings** |
| --- | --- |
| LA239 | Original Known ColRNAI with tetracycline genes |
| LA193 | Plasmid found complete on one contig |
| LA191 | Plasmid found across two contigs |
| LA189 | Original Known ColRNAI with tetracycline genes |
| LA169 | Plasmid found across two contigs |
| LA166 | Plasmid found across three contigs |
| LA139 | Plasmid found across two contigs |
| LA132 | Plasmid found across two contigs |
| LA092 | Plasmid found across three contigs |

**Supplementary S6 A) Annotated ColE1-like plasmid (pLAO84) harbouring tetracycline resistance** with Tn*1721* region (purple) and MITESen1(grey). Other prokka annotated genes shown in maroon. **B) Isolates in Illumina Laos Dataset (6) that contain this same ColE1-like plasmid**

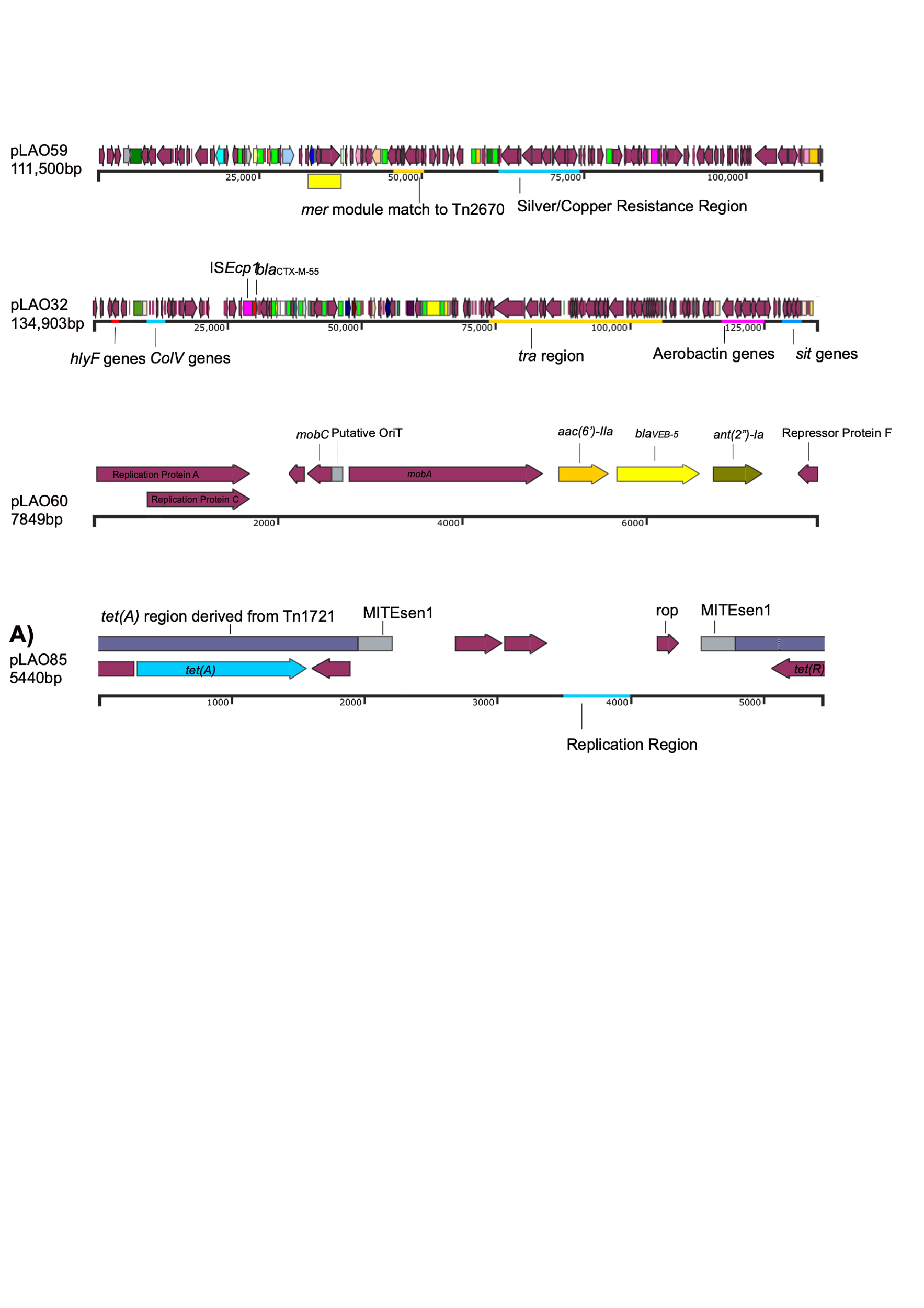

**Supplementary S7 – Annotated IncQ1 plasmid map from pLAO60 indicating the location of bla_VEB_** All maroon genes are prokka annotated genes. All other brightly coloured genes are antibiotic resistance genes.

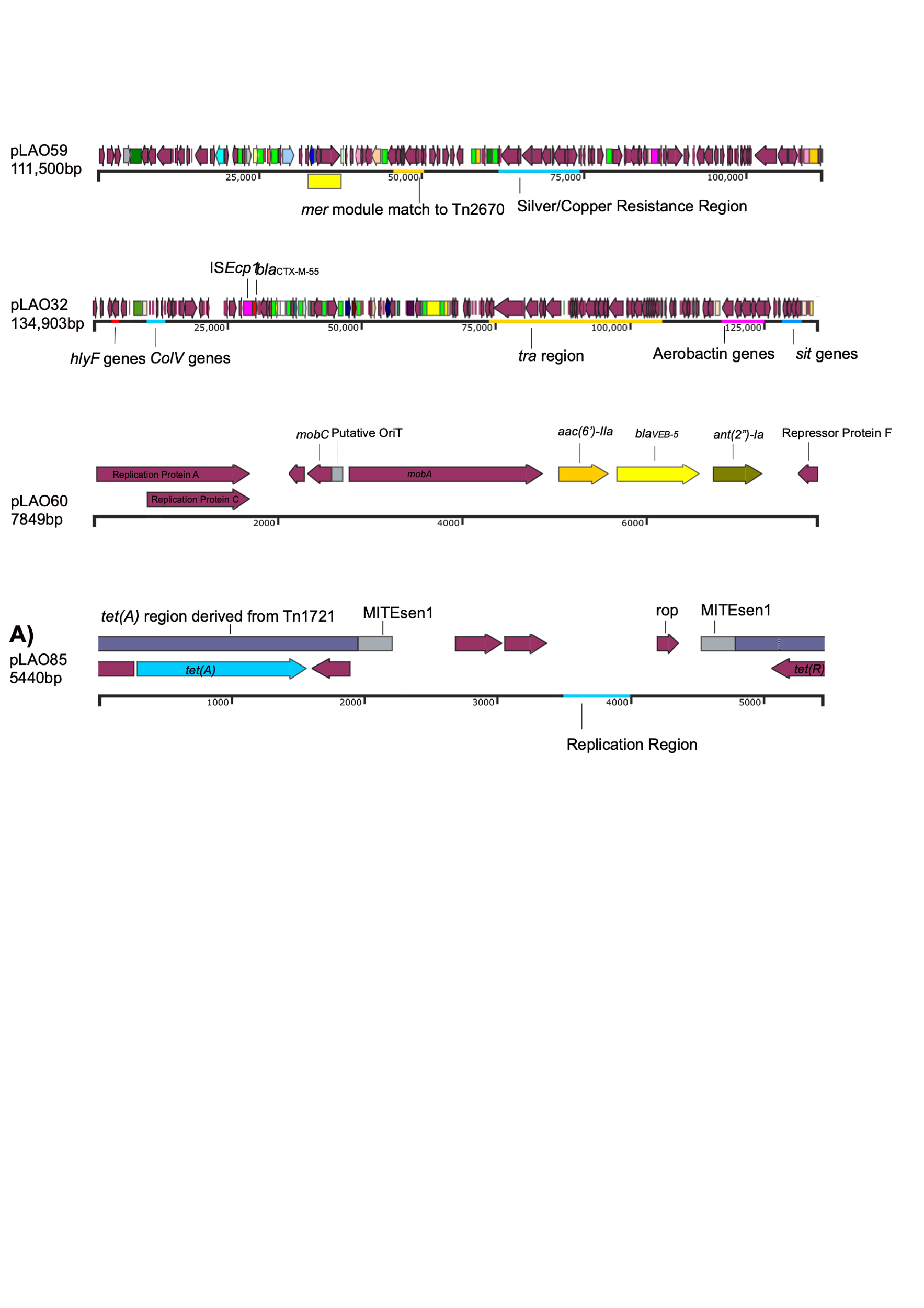

**Supplementary S8 - Annotated IncY plasmid map (pLAO59) identifying metal resistance regions and P1-like region**. All maroon genes are prokka annotated genes. All other brightly coloured genes are antibiotic resistance genes. Transposable elements (e.g. transposons, IS elements) are displayed as brightly coloured boxes. Metal resistance regions marked for copper and silver resistance(blue) and mercury resistance (orange).

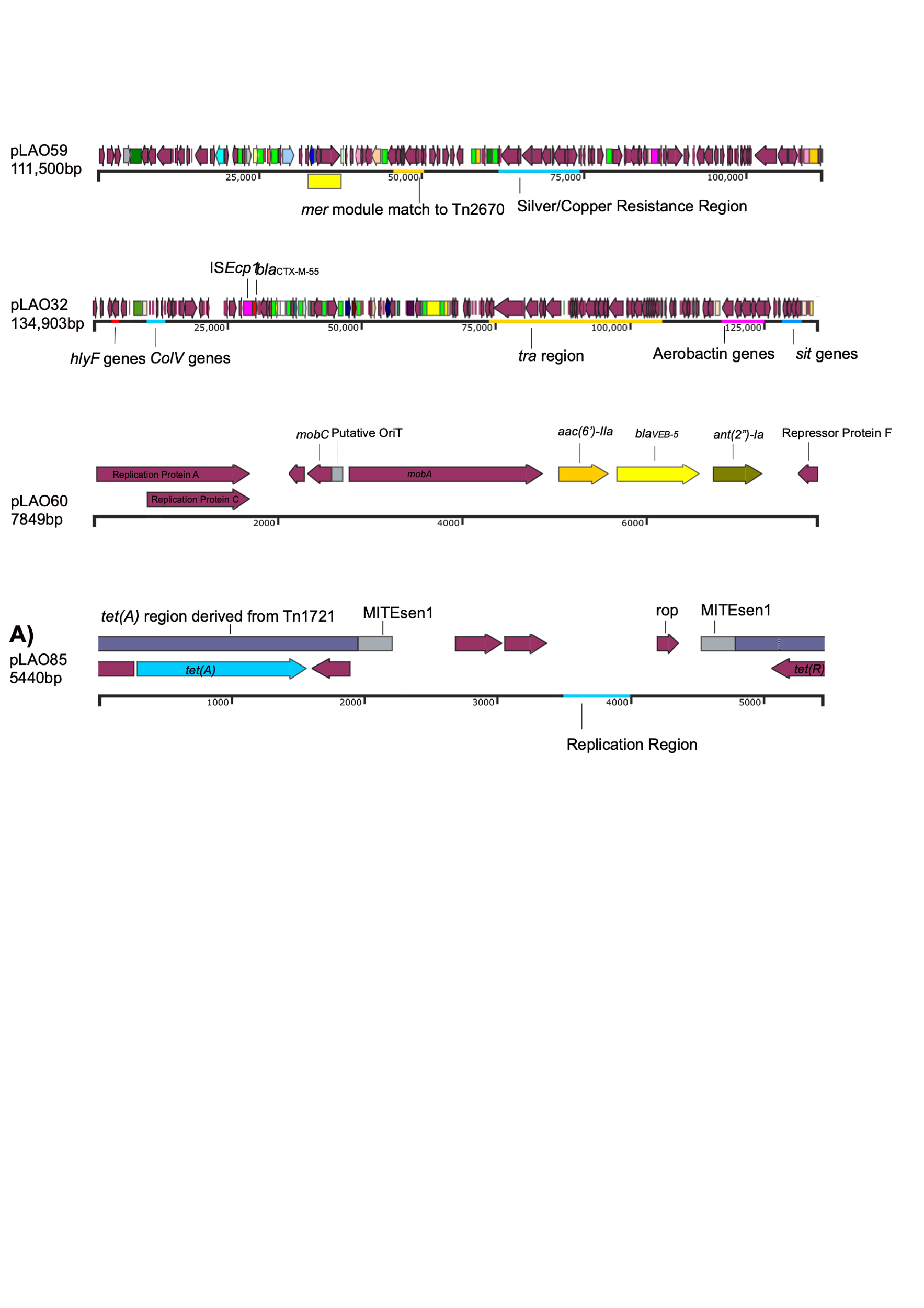

**Supplementary S9 – Annotated FII-18 plasmid map (pLAO32) showing virulence genes and *bla*_CTX-M_**. All maroon genes are prokka annotated genes. All other brightly coloured genes are antibiotic resistance genes Transposable elements (e.g. transposons, IS elements) are displayed as brightly coloured boxes. Notable IS elements are IS*26* (bright green), IS*Ecp*1 (bright pink). Transfer (*tra*) region is highlighted with the yellow outline.

**Supplementary S10** - In separate excel document

| **Plasmid** | **Query Cover (%)** | **Percentage Identity (%)** |
| --- | --- | --- |
| pLAO82 | 100 | 100 |
| pLAO37 | 100 | 100 |
| pLAO78 | 100 | 99.98 |
| pLAO86 | 47 | 92.97 |
| pLAO55 | 47 | 92.70 |
| pLAO11 | 47 | 92.70 |
| pLAO71 | 36 | 99.95 |
| pLAO61 | 36 | 99.95 |
| pLAO10 | 36 | 99.95 |
| pLAO93 | 39 | 97.24 |

**Supplementary S11 – Novel transposon Tn*7514* in other pLAO plasmids that contain complete or partial Tn*7514*.** This table shows query cover and percentage identity of Tn7514 found in these pLAO plasmids. This novel transposon has been registered with the Transposon Registry and was allocated the name Tn*7514*.

**Supplementary S12** - in separate excel file
